# Targeting *Gys1* with AAV‐SaCas9 decreases pathogenic polyglucosan bodies and neuroinflammation in Adult Polyglucosan Body and Lafora disease mouse models

**DOI:** 10.1101/2021.02.12.430952

**Authors:** Emrah Gumusgoz, Dikran R Guisso, Sahba Kasiri, Jun Wu, Matthew Dear, Brandy Verhalen, Silvia Nitschke, Sharmistha Mitra, Felix Nitschke, Berge A. Minassian

**Author notes:** **Corresponding author:** Berge Minassian, MD, Division of Neurology, Department of Pediatrics, UT Southwestern Medical Center, 5323 Harry Hines Boulevard, Dallas, TX 75390. Present affiliation: Corteva Agriscience, Johnston, IA 50131.

## Abstract

Many adult and most childhood neurological diseases have a genetic basis. CRISPR/Cas9 biotechnology holds great promise in neurological therapy, pending the clearance of major delivery, efficiency and specificity hurdles. We apply CRISPR/Cas9 genome editing in its simplest modality, namely inducing gene sequence disruption, to one adult and one pediatric disease. Adult polyglucosan body disease is a neurodegenerative disease resembling amyotrophic lateral sclerosis. Lafora disease is a severe late childhood onset progressive myoclonus epilepsy. The pathogenic insult in both is formation in the brain of glycogen with overlong branches, which precipitates and accumulates into polyglucosan bodies that drive neuroinflammation and neurodegeneration. We packaged *Staphylococcus aureus* Cas9 and a guide RNA targeting the glycogen synthase gene *Gys1* responsible for brain glycogen branch elongation in AAV9 virus, which we delivered by neonatal intracerebroventricular injection to one mouse model of adult polyglucosan body disease and two mouse models of Lafora disease. This resulted, in all three models, in editing of approximately 17% of *Gys1* alleles and a similar extent of reduction of *Gys1* mRNA across the brain. The latter led to approximately 50% reductions of GYS1 protein, of abnormal glycogen accumulation and of polyglucosan bodies, as well as corrections of neuroinflammatory markers in all three models. Our work represents proof of principle for virally-delivered CRISPR/Cas9 neurotherapeutics in an adult-onset (adult polyglucosan body) and a childhood-onset (Lafora) neurological diseases.

## Introduction

Glucose chains as short as 12 units form double helices and can precipitate, yet glycogen with up to 55,000 units is soluble. Glycogen is the product principally of two enzymes acting in concert, glycogen synthase and glycogen branching enzyme (GBE1). Glycogen synthase links glucose units through α-1,4 bonds until a certain chain length is reached, when GBE1 cleaves the terminal hexamer and reattaches it upstream through an α-1,6 bond, thus converting the chain to a fork and doubling the sites for glycogen synthase to extend. This doubling occurring with every extension generates structures that are regularly and highly branched, allowing water to permeate and retain the large molecules in solution [1]. GBE1 deficiency in mammals results in glycogen with overlong branches, which precipitates and over time accumulates into polyglucosan bodies (PBs). The precipitation is thought to be driven, like in plant starches, by the long chains winding round each other and extruding water [2, 3].

In humans, profound GBE1 deficiency leads to massive polyglucosan accumulations and early childhood death from multi-organ failure (Andersen disease) [4]. Adult polyglucosan body disease (APBD) is an allelic form of Andersen disease with lesser GBE1 deficiency (15% residual activity with the most common mutation p.Y239S). APBD patients have lesser systemic polyglucosan accumulations, and clinical disease limited to the central and peripheral nervous systems, namely a motor neuron disease with onset after age 50 and course resembling amyotrophic lateral sclerosis (ALS) [5, 6].

Two other enzymes known to regulate glycogen structure, though not understood how, are the glycogen phosphatase laforin (EPM2A) and its interacting ubiquitin E3 ligase malin (EPM2B) [7, 8]. In the absence of either, at any one time a portion of glycogen molecules appears to acquire overlong branches, leading it to precipitate and accumulate, as above, into PBs (in this context also called Lafora bodies) [3, 9]. The associated disease, Lafora disease (LD), is also neurological, characterized by teenage-onset progressive myoclonus, epilepsy and dementia, and death within 10 years of onset [9–11].

Why APBD and LD have different neurological presentations is unknown. Notwithstanding, in both the PBs drive an immune response, which at least in part underlies neurodegeneration and the neurological disease [12–20]. Since long glycogen branches are at the root of PB formation in both conditions, it was reasoned that targeting the glycogen synthase isoform expressed in the brain, *Gys1*, for downregulation may represent a therapeutic modality for both diseases. APBD (*Gbe1*^*Y239S*^) and LD (*Epm2a*^*−/−*^ and *Epm2b*^*−/−*^) mouse models replicate their corresponding diseases [21–23]. Crossing these mice with mice deficient of GYS1 activity dramatically reduced PBs and neuroimmune markers and rescued behavioral phenotypes [12, 13, 17, 19, 20, 24–26], confirming the role of PB accumulation in the pathogeneses of these diseases, and supporting the targeting of GYS1 as a potential, shared, therapeutic approach.

CRISPR/Cas9 genome editing is widely expected to revolutionize neurology [27–29]. Among the hurdles to be overcome are achieving wide distribution of the enzyme/guide complex across the brain, and accurate target editing. The latter is not a concern when the goal is not actually to edit, but to delete a gene function. In the present work we show that AAV9-delivered *Gys1*-directed CRISPR/Cas9 significantly reduces PB accumulations and immune markers in APBD and LD mouse models. Our results represent proof of principle for GYS1-targeting CRISPR/Cas9 as a potential therapy for both these fatal diseases.

## Methods

### Study design

We hypothesized that CRISPR/Cas9–mediated disruption of the *Gys1* gene in *Gbe1*^*Y239S*^, *Epm2a*^*−/−*^ and *Epm2b*^*−/−*^ mice would improve the neuropathology of these models. Following packaging in adeno-associated virus 9 (AAV9), we administered *Gys1-*targeting CRISPR/Cas9 at postnatal day 2 by intracerebroventricular (ICV) injection, sacrificed the mice at 3 months of age, and studied the effect of the treatment on PB quantity and neuroinflammation markers.

### Plasmid construction and viral packaging

We selected multiple *Gys1* targeting candidate guide RNAs (sgRNAs) using the CRISPR Design Tool software [30], which we custom-synthesized to include 5’-end guanines for efficient expression from the U6 promoter. These were cloned into a published *Staphylococcus aureus* Cas9 (SaCas9)-expressing AAV vector [31], replacing that vector’s SaCas9 CMV promoter with the small synthetic ubiquitously expressing JeTI promoter for more efficient viral packaging [32], with final arrangement as in Fig. 1. Candidate plasmids were transfected into mouse N2A cells, with genomic DNA extracted 48 h post-transfection for the Inference of CRISPR Edits assay (ICE), as published [https://tinyurl.com/y43gtzux], to identify the sgRNA causing the most indel formation (final plasmid ICE primer and sgRNA sequence in Table 1). The final plasmid was packaged into AAV9 at the University of North Carolina (UNC) Vector Core facility, as described [33].

**Table 1.**
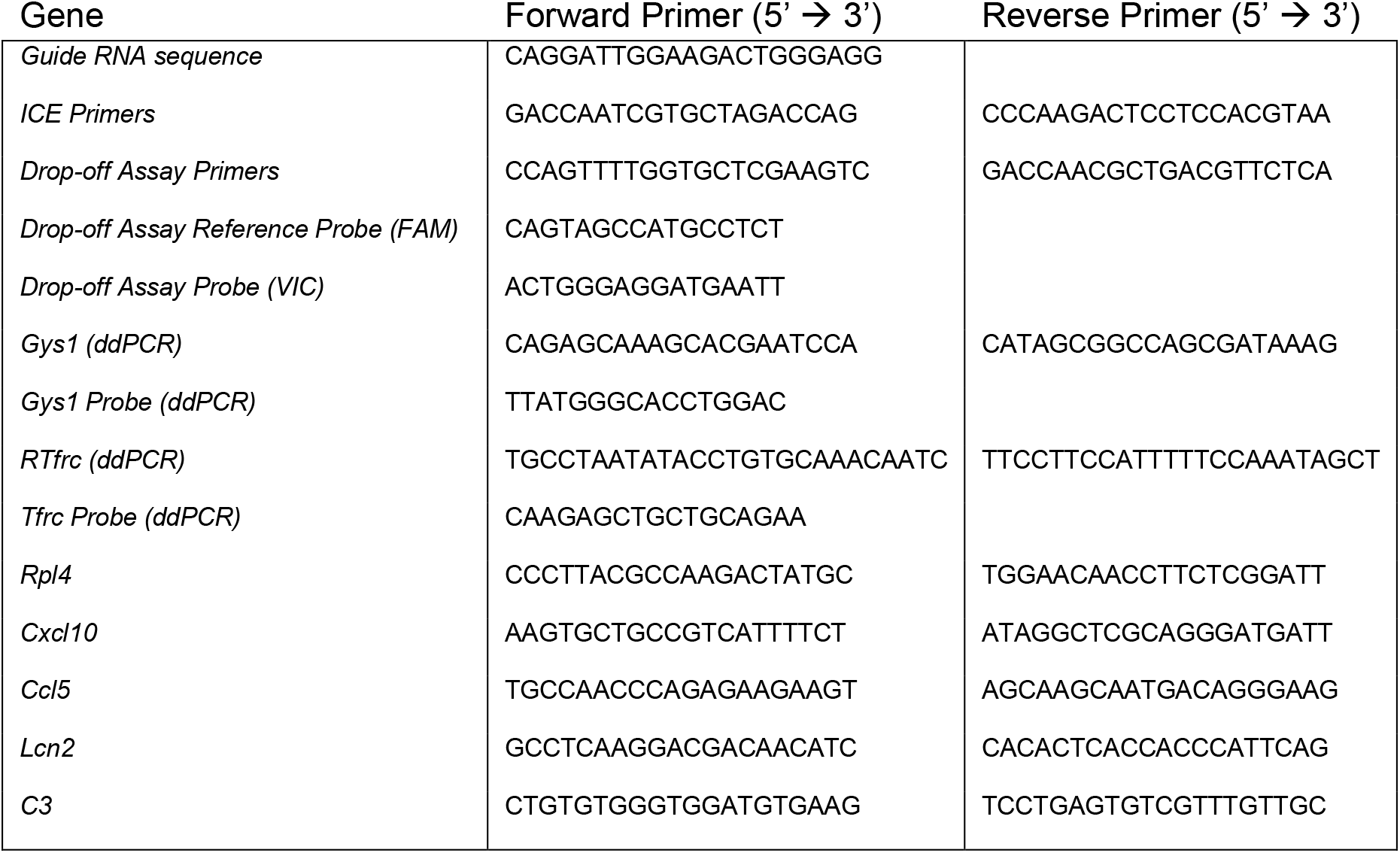
Primer and probe sequences used in this study

**Fig. 1:**
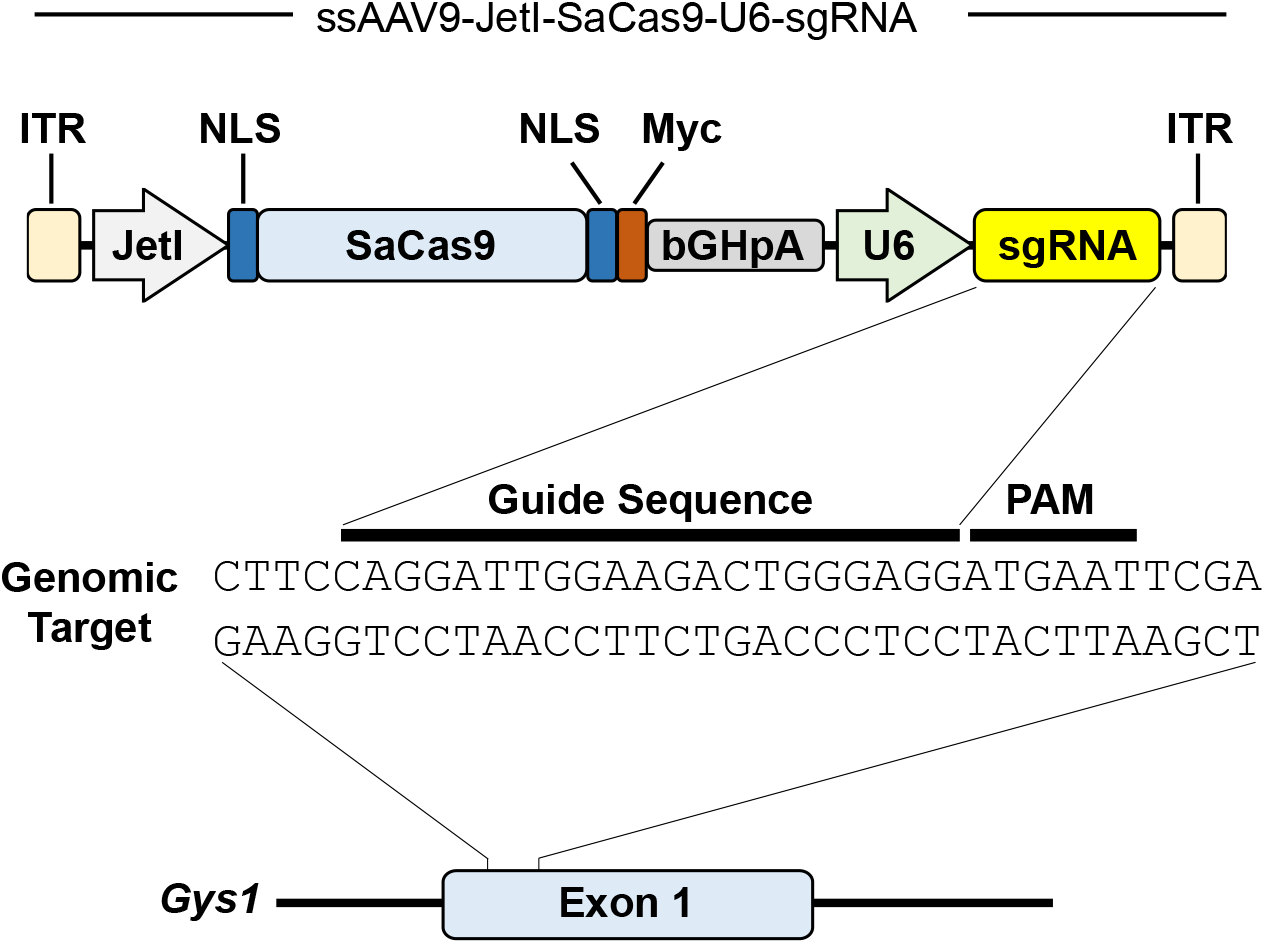
AAV-SaCas9 targeting murine *Gys1*. Schematic representation of AAV vector, guide RNA sequence, and *Gys1* - Exon1 target site. ITR, inverted terminal repeats; Myc, c-myc; NLS, nuclear localization signal sequence; bGH-pA, bovine growth hormone polyadenylation signal; PAM, protospacer adjacent motif.

### Mice

APBD (*Gbe1*^*Y239S*^) and LD (*Epm2a*^*−/−*^ and *Epm2b*^*−/−*^) mouse models were described previously [21–23]. Of relevance to the interpretation of some of the results (see penultimate paragraph in the discussion section) that while APBD is ‘milder’ than LD (later age of onset and slower progression) in humans, the reverse is true in mouse where APBD mice die by approximately 10 months of age, while LD mice only begin to have mild behavioural phenotypes at that age [21–23]. Both sexes were used in approximately equal proportion in all experiments. All procedures were carried out according to NIH guidelines and the animal care committee regulations at the University of Texas Southwestern Medical Center. 7×10^11^ vector genomes (as 5 μl solution) or 5 μl PBS were injected ICV, as described [34]. Mice were sacrificed by cervical dislocation, and brain harvested and cut into two hemispheres, one fixed in formalin for paraffin embedding and histo- and immunohisto-chemistry, the other snap-frozen in liquid nitrogen, ground into powder on the same by mortar and pestle, and aliquoted into 30 - 40 mg powder in screwcap tubes for the genetic and biochemical experiments below.

### Cas9 immunohistochemistry

Sections (5 μm) were mounted on glass slides, de-paraffinized and rehydrated by processing through xylenes and decreasing concentrations of ethanol in water, and subjected to antigen retrieval using citrate buffer pH 6.0 (Sigma). Endogenous peroxidase activity was blocked for 10 min with BLOXALL solution (Vector labs). Sections were incubated with rabbit anti-CRISPR-Cas9 antibody (1:50, Abcam) diluted in normal horse serum overnight at 4°C, then successively incubated with Amplifier Antibody and ImmPRESS Polymer Reagent (Vector labs) and the ImmPACT DAB EqV working solution (Vector labs) until desired stain intensity.

### *Gys1* indel and expression quantifications by droplet digital PCR

Drop-off assay (QX200 Droplet Digital PCR system, Bio-Rad Laboratories) was used to quantify the indels generated by AAV-SaCas9 per manufacturer’s instructions. Genomic DNA was extracted using PureLink Genomic DNA Mini Kit (Invitrogen). 40X Taqman SNP genotyping assay and custom designed primer-probe set (Life Technologies, Table 1) was used to detect indels. Briefly, the primer set amplifies the sequence that includes the predicted cut site, a FAM-labeled reference probe distant from the site of potential non-homologous end joining site counts all copies of the amplicon, and a VIC-labeled drop-off probe binds the predicted cut site. The reaction mix consisted of 10 ml of 2x ddPCR SuperMix for Probes (Bio-Rad Laboratories), 0.5 ml of the 40X assay, 9.5 ml nuclease-free water and 1.0 ml gDNA. Cycling conditions were 95°C for 10 min, 45 cycles of 94°C for 30 sec and 58°C for 1 min, 98°C for 10 minutes and finally a 10°C hold on a Life Technologies Veriti thermal cycler. The loss of VIC signal (while maintaining FAM signal) was used as an indel indicator. The resulting data was quantified by QuantaSoft v1.4 (Bio-Rad).

RNA was extracted using TriZol (Invitrogen) and purified using PureLink RNA Mini Kit (Invitrogen) following manufacturer instruction. cDNA was generated using the iScript Reverse Transcription SuperMix kit (Bio-Rad Laboratories). *Gys1* RNA was quantified using the Bio-Rad QX200 Droplet Digital PCR (ddPCR) system as instructed by the manufacturer. *Tfrc* was used as reference gene. Custom designed TaqMan primers and probes (Thermo Fisher Scientific) were used for both *Gys1* and *Tfrc*. ddPCR reactions were assembled using standard protocols. The 20 μL reaction mix consisted of 10 μL 2x ddPCR SuperMix for Probes (Bio-Rad Laboratories), 1 μL *Gys1* assay mix (FAM-labeled), 1 μL reference *Tfrc* assay mix (VIC-labeled), 3 μL of cDNA, and 5 μL nuclease-free water. Cycling conditions were 95 °C for 10 min, 45 cycles of 94 °C for 30 sec and 60 °C for 1 min, and 98 °C for 10 min. PCR and data analysis equipment were the same as above. Results were expressed as ratio of *Gys1*/*Tfrc*. No-template controls were run in parallel with study samples. Primer/probe sets are in Table 1.

### SaCas9 and GYS1 western blots

Tissue lysate was obtained by homogenizing frozen ground brain tissue using ice-cold RIPA lysis buffer (NaCl 150 mM, NP-40 1%, Sodium deoxycholate 0.5%, SDS 0.1%, Tris HCl 50 mM pH 8.0) containing protease inhibitors (1mM PMSF, 5 μg/mL Leupetin, 10 μg/mLPepstatin, 20KIU/mL Aprotinin, 50mM NaF). Lysates were centrifuged at 15,000 x g for 5 min at 4 °C and supernatants collected. Protein concentration was measured using the Bradford assay reagent (ThermoScientific). Serial dilution of albumin standard (ThermoScientific) generated the standard curve for protein quantification. Equal amounts of whole protein from each sample was subjected to SDS-PAGE using the TGX Stain-Free FastCast Acrylamide kit (BioRad). Protein bands were transferred to polyvinylidene difluoride (PVDF) membrane (Millipore) overnight at 4°C. Primary antibodies for SaCas9 (1:1000, abcam) and for GYS1 (1:1000, Cell Signaling) were used. Protein dansitometry was performed using Image Lab software (Bio-Rad Laboratories). Intensity of each protein band was normalized to the intensity of its corresponding whole protein lane image obtained on the same membrane.

### PASD staining and LB quantitation

Paraffin-embedded brain tissue was sectioned and stained using the periodic acid-Schiff diastase (PASD) method as described previously [13, 20]. Stained slides were scanned using the Hamamatsu Nanozoomer 2.0 HT digital slide scanner (40 x objective), and the % area of hippocampus covered by LBs was quantified using ImageJ [35] and iLastik (v. 1.3.3) [36].

### Quantification of degradation-resistant glycogen

Aliquot of frozen ground brain was left out to thaw at room temperature for 1 hour to allow soluble glycogen to degrade, leaving behind degradation-resistant, i.e. PB, glycogen [37]. The sample was then boiled in 30% KOH and glycogen precipitated in 70% ethanol with 57 mM sodium sulfate. Three further rounds of centrifugation and suspension in 67% ethanol with 15 mM LiCl followed after which the pellet was resuspended in 100 mM sodium acetate pH 4.5 and digested with amyloglucosidase (Megazyme) in a 55°C oven for 1 hour along with digestion of blank controls. Glucose amount was determined in triplicate for each sample using a modified version of Lowry and Passonneau [38]. Briefly, samples and glucose standards were mixed with 170 μL of G6PDH reaction mix (200 mM tricine buffer/10 mM MgCl_2_ pH 8, 0.66 mM NADP; 1 mM of ATP; 0.5 units of G6PDH). Absorbance at 340 nm was acquired before (Abs1) and after (Abs2) addition of hexokinase (0.6 units in 4 μl of 200 mM Tricine/KOH, pH 8, 10 mM MgCl_2_). (Abs2 – Abs1) of glucose standards was used to derive glucose quantity in samples. Following normalization to fresh weight, glucose quantity represents degradation-resistant glycogen [37].

### Quantitative real-time PCR

Immune system related genes *Cxcl10, Ccl5, Lcn2 and C3* were quantified by quantitative real-time PCR (qRT-PCR) using the QuantStudio 7 Pro System thermo-cycler (ThermoFischer Scientific) and SYBR Green Master Mix (Bio-Rad Laboratories). Data are shown as fold change relative to control samples using the ΔΔCq method with *Rpl4* as an internal control gene. Primers are listed in Table 1.

### Statistical analysis

Student’s unpaired t-test was used to compare single means. Data were analyzed and graphed using the GraphPad Prism software (v. 8.0.2; GraphPad Software). For all comparisons, statistical significance was set at *p* < 0.05. Asterisks denote level of significance based on *p* value: \**p* < 0.05, \*\**p* < 0.01, \*\*\**p* < 0.001, and \*\*\*\**p* <0.0001.

## Results

### Extents of SaCas9 distribution and activity

For a visual assessment of the pattern and extent of virally delivered SaCas9 distribution, we performed immunohistochemistry in one of our animal models (*Gbe1*^*Y239S*^) using an anti-SaCas9 antibody. Large numbers of cells expressed the enzyme in brain regions surrounding the cerebral ventricles, namely the cortex, hippocampus and thalamus/hypothalamus. Expressing cells became sparser the further from the ventricles to almost none in the cerebellum (Fig. 2).

**Fig. 2:**
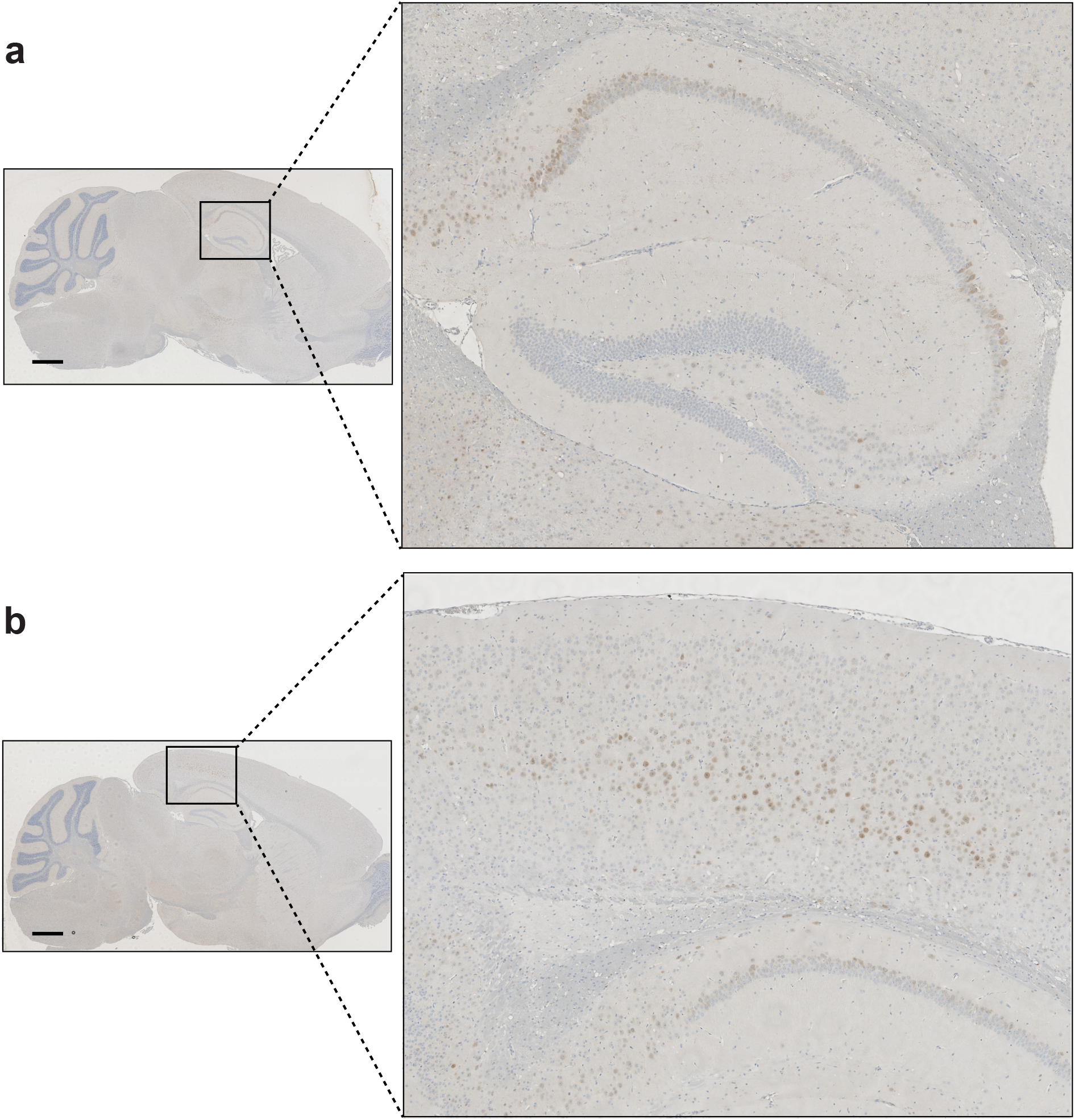
AAV-SaCas9 virus distribution as detected by SaCas9 immunohistochemistry. Representative images show SaCas9 expressing cells in hippocampus (a) and cortex (b). Scale bar is 1mm.

To quantify the number of *Gys1* alleles disrupted by the *Gys1*-targeted SaCas9, we used ddPCR and obtained the percentage of total *Gys1* with indel formations in extracts from aliquots of frozen powdered whole hemispheres. In all three mouse models (*Gbe1*^*Y239S*^, *Epm2a*^*−/−*^ and *Epm2b*^*−/−*^) approximately 17% of *Gys1* alleles had been edited (Figs. 3a, 4a, 5a).

**Fig. 3:**
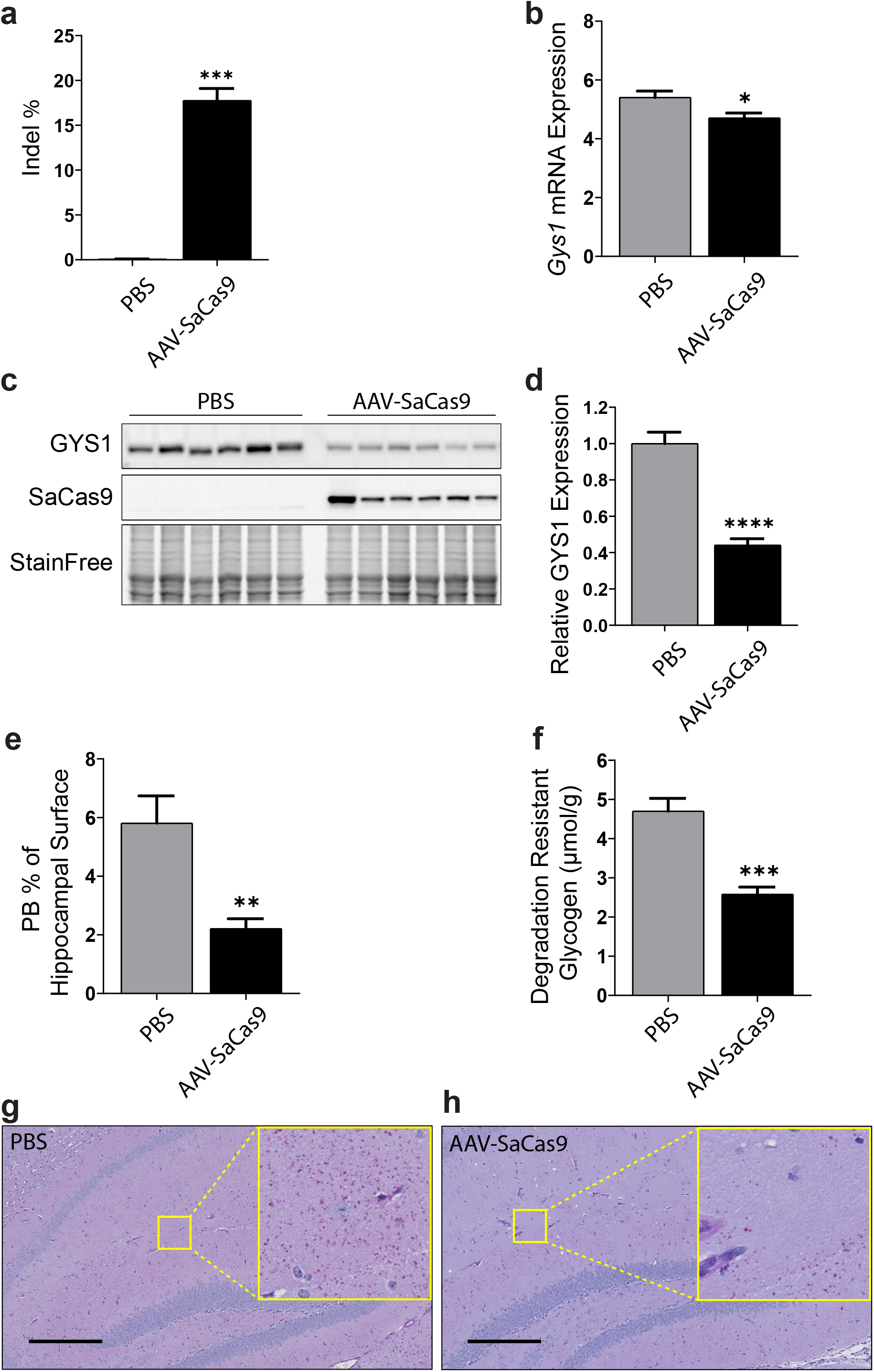
*Gys1* targeting AAV-SaCas9 disrupts the *Gys1* gene, reduces Gys1 mRNA and protein levels, and decreases insoluble glycogen and PB accumulation in brains of the *Gbe1*^*Y239S*^ APBD mouse model. Neonatal mice (P2) were injected with PBS (N = 12 for each experiment) or AAV-SaCas9 (N = 12 for each experiment), and mice were sacrificed at 3 months for brain tissue analysis. Indel percentage (a) and Gys1 mRNA level (b) were measured by ddPCR. Representative brain GYS1 western blots with stain-free gel as loading control (c). Quantification of GYS1 western blots normalized to stain-free gel shown in (d). Polyglucosan body (PB) quantification in the hippocampus (e) and degradation resistant glycogen content (f). Representative micrographs of PASD stained hippocampus of PBS (g) vs AAV-SaCas9 (h) treated mouse. Scale bar is 300 μm. All data are presented as mean ± SEM. Significance levels are indicated as *, p < 0.05; **, p < 0.01; ***, p < 0.001; ****, and p < 0.0001.

**Fig. 4:**
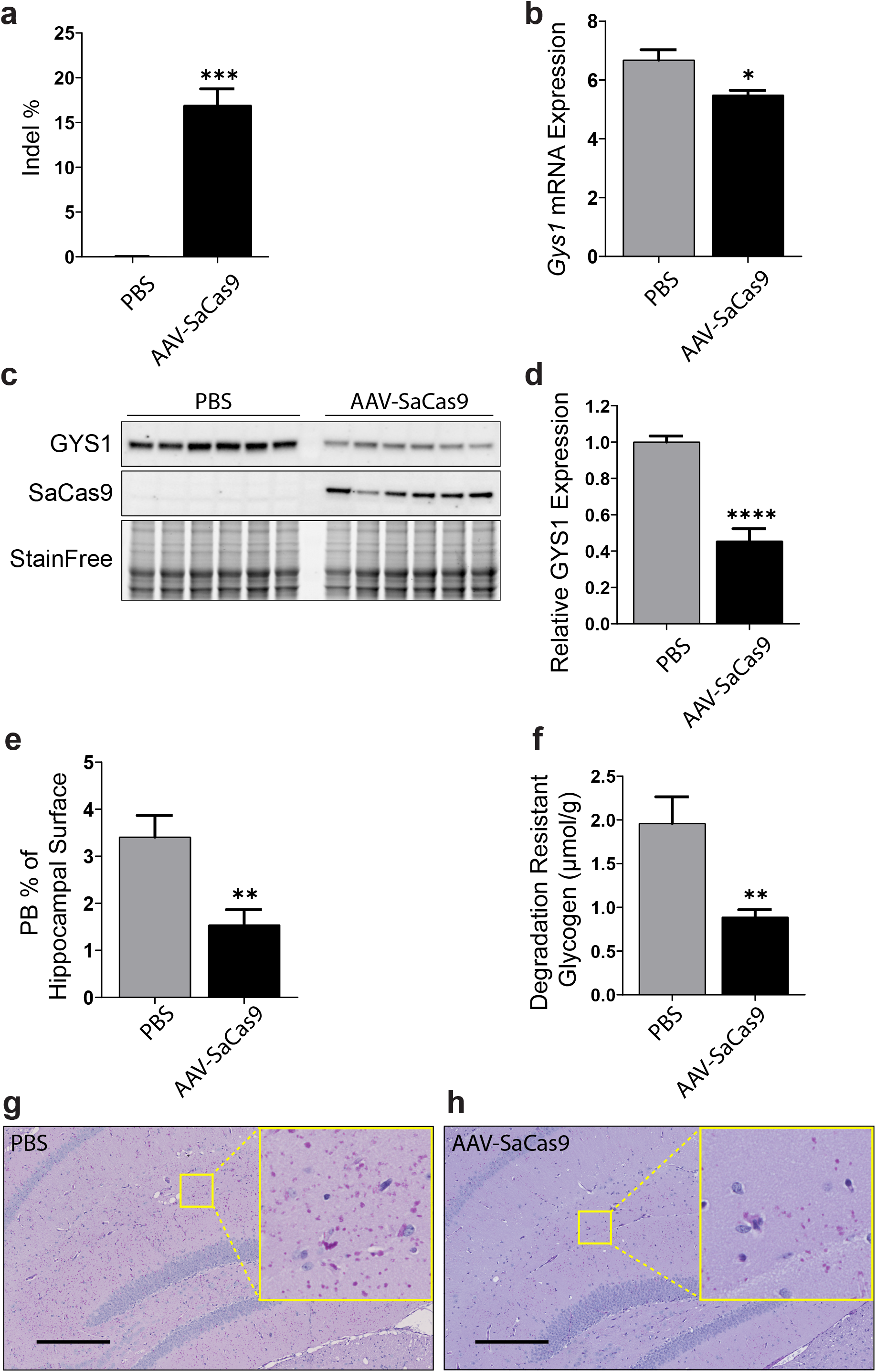
*Gys1* targeting AAV-SaCas9 disrupts the *Gys1* gene, reduces Gys1 mRNA and protein levels, and decreases insoluble glycogen and PB accumulation in brains of the *Epm2a*^*−/−*^ LD mouse model. Neonatal mice (P2) were injected with PBS (N = 13 for each experiment) or AAV-SaCas9 (N = 8 in WB quantification and N = 10 for other experiments), and mice were sacrificed at 3 months for brain tissue analysis. Indel percentage (a) and Gys1 mRNA level (b) were measured by ddPCR. Representative brain GYS1 western blots with stain-free gel as loading control (c). Quantification of GYS1 western blots normalized to stain-free gel shown in (d). Polyglucosan body (PB) quantification in the hippocampus (e) and degradation resistant glycogen content (f). Representative micrographs of PASD stained hippocampus of PBS (g) vs AAV-SaCas9 (h) treated mouse. Scale bar is 50 μm. All data are presented as mean ± SEM. Significance levels are indicated as *, p < 0.05; **, p < 0.01; ***, p < 0.001; ****, and p < 0.0001.

**Fig. 5:**
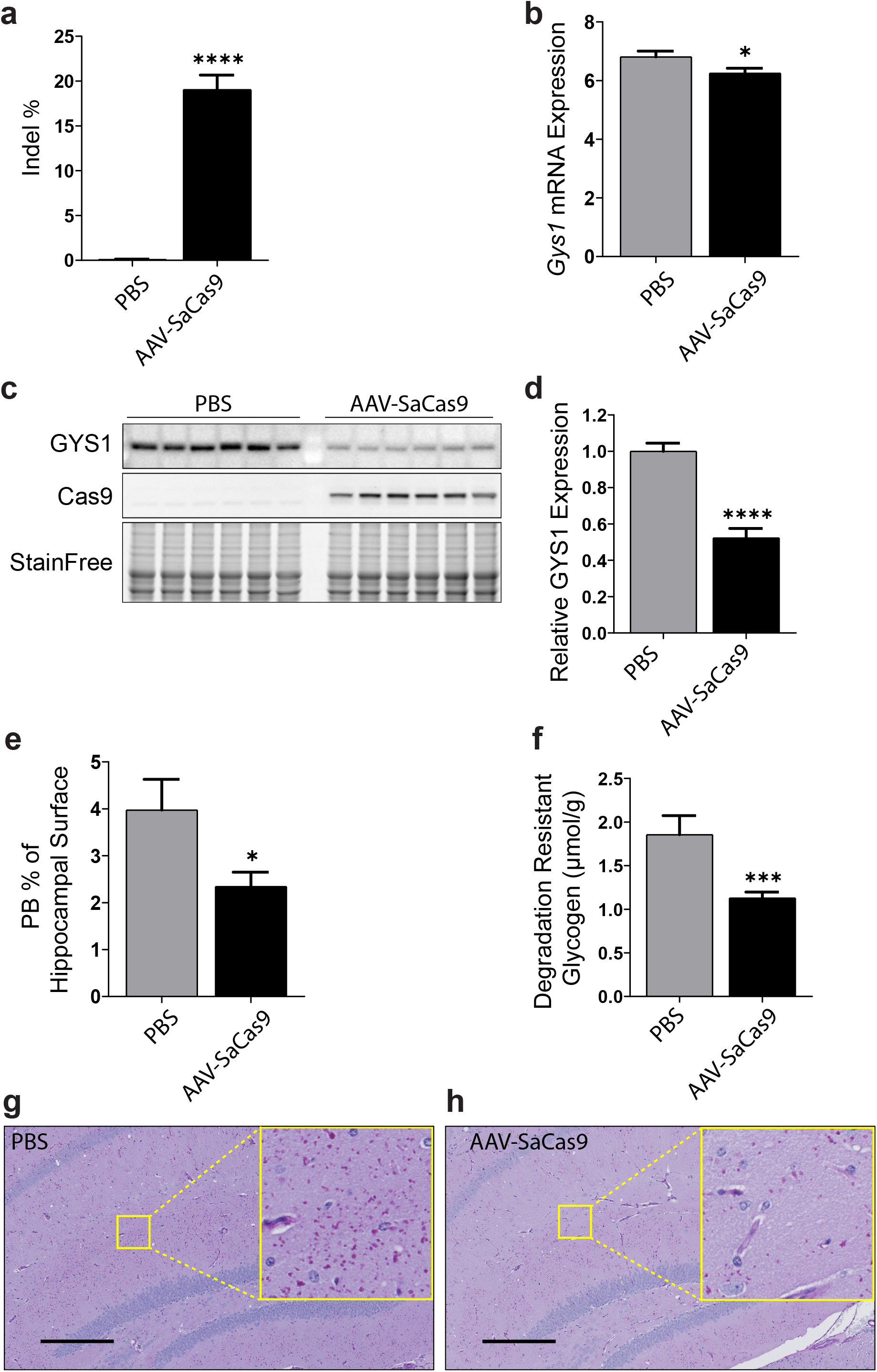
*Gys1* targeting AAV-SaCas9 disrupts the *Gys1* gene, reduces Gys1 mRNA and protein levels, and decreases insoluble glycogen and PB accumulation in brains of the *Epm2b*^*−/−*^ LD mouse model. Neonatal mice (P2) were injected with PBS (N = 10 for each experiment) or AAV-SaCas9 (N = 14 for each experiment) and mice were sacrificed at 3 months for brain tissue analysis. Indel percentage (a) and Gys1 mRNA level (b) were measured by ddPCR. Representative brain GYS1 Western blots with stain-free gel as loading control (c). Quantification of GYS1 Western blots normalized to stain-free gel shown in (d). Polyglucosan body (PB) quantification in the hippocampus (e) and degradation resistant glycogen content (f). Representative micrographs of PASD stained hippocampus of PBS (g) vs AAV-SaCas9 (h) treated mouse. Scale bar is 50 μm. All data are presented as mean ± SEM. Significance levels are indicated as *, p < 0.05; **, p < 0.01; ***, p < 0.001; ****, and p < 0.0001.

### Effects on *Gys1* mRNA and protein

To measure the extent of reduction of whole hemisphere *Gys1* mRNA, we again used ddPCR on whole hemisphere extracts and found an approximately 15% reduction in the AAV-Cas9-treated mice of all three genotypes (Figs. 3b, 4b, 5b), a value not dissimilar to the extent of *Gys1* DNA rearrangement. Reduction of GYS1 protein measured by quantitative western blotting on whole hemisphere extract was by approximately 50% in all three disease models (Figs. 3c, d; 4c, d; 5c, d), unexpectedly higher than the mRNA reduction.

### Effects on hippocampal LBs and whole hemisphere degradation-resistant glycogen

In all three disease models, PBs are more or less evenly distributed across the brain [21–23]. In most studies, PB measures are obtained in the hippocampus, because this region’s characteristic structure permits demarcation of a uniform tissue surface area for reliable measurements across experimental animals [13, 17, 26]. Hippocampal PB loads measured as % PASD signal per hippocampal area were reduced by 40 to 60% in the AAV-Cas9-treated mice in the three models (Figs. 3e, g, h; 4e, g, h; 5e, g, h), a reduction that is in the range of GYS1 protein diminution.

In oxygen and glucose deprived states, normal brain glycogen is very swiftly digested to generate glucose in an attempt to protect the brain [39, 40]. This is so rapid that within minutes of sacrifice normal murine brain glycogen all but disappears [37]. In the APBD and LD mouse models, brain glycogen is a mix of normally branched soluble and consumable glycogen and the ever-accumulating abnormally branched precipitated and digestion-resistant polyglucosans (PBs). Within 60 minutes from sacrifice, the former disappear while the latter remain and can be biochemically quantified as a measure of PBs [3, 37, 41]. This degradation-resistant glycogen was reduced by 40 to 60% in whole hemisphere extracts in AAV-Cas9 treated mice in the three genotypes (Figs. 3f, 4f, 5f), a value similar to the reduction in PBs as quantified histochemically.

### Effects on PB-associated immune activation

Recent studies in LD mouse models revealed an important inflammatory component in disease pathogenesis. In advanced disease (over 11 months of age) immunohistochemical studies showed activation and proliferation of astroglia and microglia. Transcriptomic analyses showed that 94% of genes upregulated in the disease encode proteins of inflammatory and immune system pathways [14]. Finally, downregulation of GYS1 activity by transgenic or other means resulting in PB reductions partially corrected these abnormalities [15–20]. Of the several hundred immune system genes shown to be dysregulated in the transcriptomic studies, a subset (less than 10) was tested and confirmed by qRT-PCR, of which several were shown to already be overexpressed at earlier stages of the disease [14, 20]. To our knowledge similar transcriptomic, and generally RNA based studies of immune markers, have not been performed in APBD mice, although the latter are known to exhibit astrogliosis and microgliosis in immunohistochemical studies [13].

We selected the four most studied immune marker genes and quantified their expression in the current cohorts. At the age of sacrifice (3 months) three (*Cxcl10, Lcn2,* and *C3*) of the four were upregulated in the APBD mice and all three were corrected with the *Gys1*-targeting AAV-Cas9 (Fig. 6a). Two genes (*Cxcl10* and *Ccl5*) were upregulated in the *Epm2a*^*−/−*^ mice, and corrected with the treatment (Fig. 6b). One gene (*Cxcl10*) was upregulated in the *Epm2b*^*−/−*^ mice, and corrected.

**Fig. 6:**
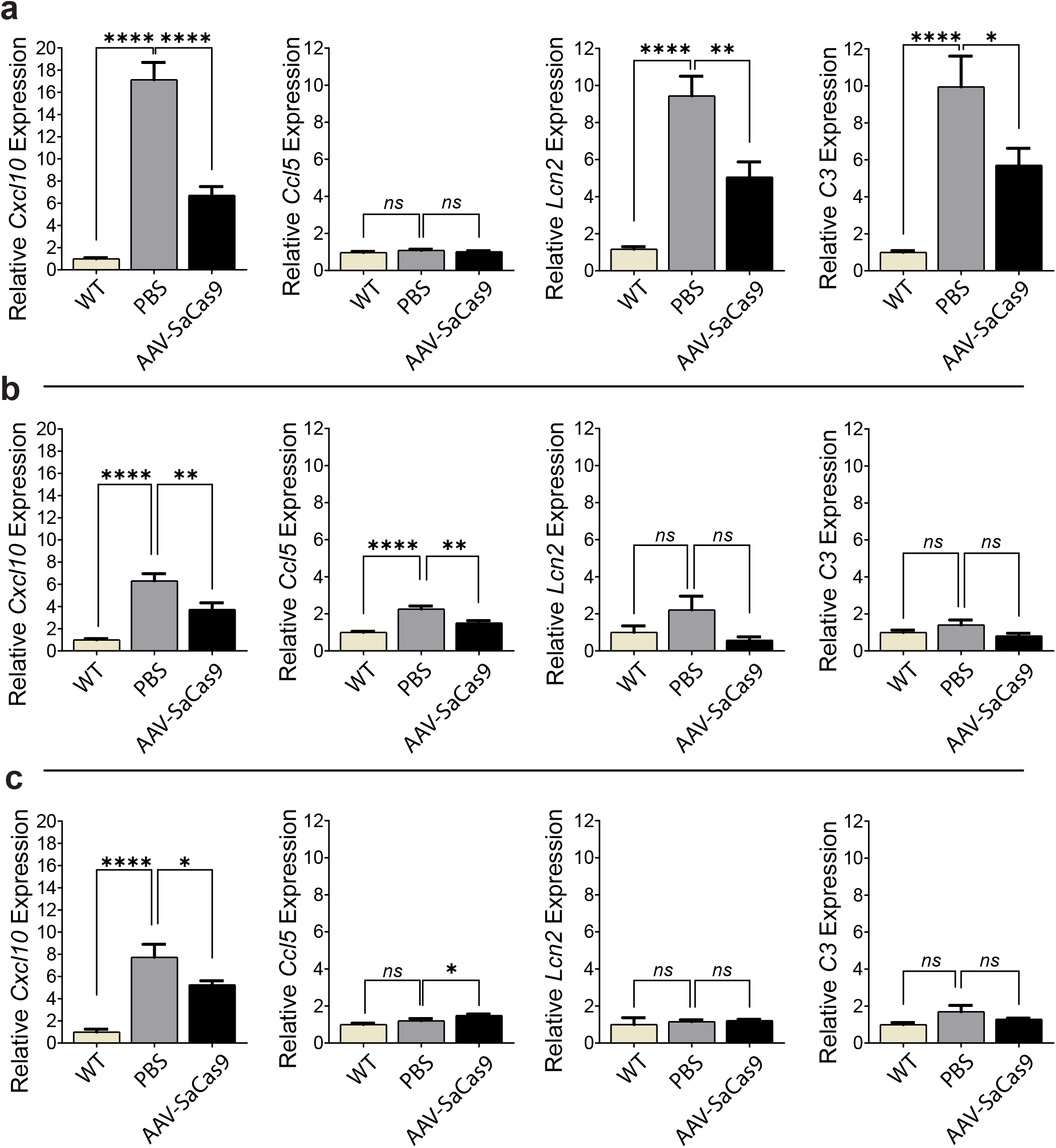
Effect of AAV-SaCas9 treatment on polyglucosan body associated immune system activation. Neonatal mice (P2) were injected with PBS or AAV-SaCas9 and mice were sacrificed at 3 months for brain tissue analysis. WT indicates the wild type control group. For each panel, from left to right, *Cxcl10, Ccl5, Lcn2* and *C3* were used as neuroinflammation markers, and relative mRNA expression levels were analyzed by qRT-PCR for *Gbe1*^*Y239S*^ (a), *Epm2a*^*−/−*^ (b) and *Epm2b*^*−/−*^ (c) mice. In panel (a); WT (N = 13), PBS (N = 12) and AAV-SaCas9 (N = 12). In panel (b); WT (N = 8), PBS (N = 13), and AAV-SaCas9 (N = 10). In panel (c); WT (N = 11), PBS (N = 10), and AAV-SaCas9 (N = 14). All data are presented as mean ± SEM. Significance levels are indicated as *, p < 0.05; **, p < 0.01; ***, p < 0.001, ****, and p < 0.0001.

## Discussion

Given its complexity, the brain utilizes the lion’s share of the genome for its development and function. As such, thousands of single gene defects present with neurological problems in childhood, the corollary of which is that child neurology in large part consists of numerous rare monogenic diseases. A much smaller number of single gene defect disorders presents in adulthood, but many adult neurological and neuropsychiatric diseases are still genetic, albeit complex, including common forms of Alzheimer’s disease, schizophrenia, Parkinson’s disease, ALS, etc. As such, gene-based therapies are a definite part of the future of clinical neurosciences.

At present, CRISPR/Cas based technologies are the most promising in editing the genome back to health. Two of the major hurdles to applicability are transport of the molecular machinery into vast swathes of brain cells, and ensuring accurate editing. These two difficulties are interrelated as the latter requires inclusion of components that makes the effective cargo that much more complex and larger. The simplest form of editing is gene disruption, which is achievable with only two components, a specific guide RNA and a Cas enzyme. The diseases studied here represented an opportunity to test this basic possibility. The AAV9 virus accommodated the comparatively small SaCas9 enzyme and a *Gys1*-targeting guide in a single plasmid that was delivered and was functional on 17% of *Gys1* alleles across the brain.

The above ~17% disruption was achieved by ICV injection in neonatal mice, of all known situations the most permissive given the yet immature nature of intercompartmental barriers at this age [42–44]. Treatments at older ages through any route (ICV, intrathecal, intra-cisterna magna, or intravenous) are expected to be less widespread and effective, and will require enhanced viral or other delivery systems.

The most commonly used streptococcal Cas9 (~4 kb) does package in AAV9 but cannot accommodate the guide sequence, thus requiring separate viruses delivering the two necessary components to the same cells, which greatly reduces editing efficiency. The SaCas9 ortholog used here (~3.2 kb) does accommodate the guide within one virus [45]. The extent of viral distribution is greatly enhanced with self-complementary vector designs, where two copies of the cargo sequences are incorporated [46, 47]. This of course could not be applied here, given the size even of SaCas9. Future developments in obtaining much smaller Cas enzymes or larger packaging capacity would improve self-complementation-based spread of effective delivery.

The JeTI promoter used in our construct to drive SaCas9 expression is a short artificial promoter designed to make room to fit larger genes within AAV9’s packaging capacity [48, 49]. Its expression potency is low compared to standard gene therapy promoters, which in this case may be an advantage. Cas9 is needed to act twice in each cell, once on each autosome target. Any activity beyond that would be off-target and potentially detrimental, and as such weak promoters are likely preferable.

In our study, the extent of *Gys1* mRNA reduction corresponded to that of *Gys1* gene editing (15-17%). Surprisingly, the degree of GYS1 protein reduction was substantially higher (~50%). We observed a similar greater GYS1 protein than mRNA reduction in our previous work where we knocked out the *Gys1* gene through a conditional transgenic approach [20]. Generally, disconnect between mRNA versus protein relative quantities is not surprising and well-documented, with a multiplicity of post-transcriptional mechanisms invoked and under study [50]. In the present case the disconnect is favourable: treatments for APBD and LD with finite impacts on *GYS1* mRNA would have a greater impact on the ultimate enzyme target.

The recent recognition of the inflammatory component in the pathogenesis of LD opened novel anti-inflammatory therapeutic possibilities, which are presently being studied, while more permanent root cause-based therapies are developed [14–16, 51]. A side result of our study is the identification of similar transcription-level immune abnormalities, and their correction, in APBD. These data supplement our recent immunohistochemical findings of astrogliosis and microgliosis in APBD and their correction following transgenic targeting of the *Gys1* gene [13]. Collectively, these results suggest that the immune pathology, and its response to GYS1 and PB lowering therapies, may be shared across polyglucosan storage diseases. In fact, the immune dysregulation appears to be greater in the APBD mouse model than in the LD models, with more immune markers affected at the early age of three months. This greater immune disturbance correlates with the larger number of PBs and the greater accumulation of digestion-resistant glycogen in the APBD versus LD models (compare panels e and f across Figs. 3 to 5), further supporting a relationship between PB accumulation and immunopathology.

In conclusion, it is possible to achieve correction of pathological bases of neurological diseases of the brain through AAV9-delivered Cas9 genome editing. Immunopathology may be shared across brain polyglucosan storage diseases. A single therapy, targeting *GYS1*, may be possible for multiple glycogen storage diseases of the brain.

## Acknowledgments

This work was funded by the National Institutes of Health under award P01NS097197. B.A.M. holds the University of Texas Southwestern Jimmy Elizabeth Westcott Chair in Pediatric Neurology. We thank Drs. Feng Zhang (Massachusetts Institute of Technology and Harvard University) and Steven Gray (University of Texas Southwestern) and team members for their help in vector design, construction and packaging.

